# Pyrolyzed substrates induce aromatic compound metabolism in the post-fire fungus, *Pyronema domesticum*

**DOI:** 10.1101/2021.03.15.435558

**Authors:** Monika S. Fischer, Frances Grace Stark, Timothy D. Berry, Nayela Zeba, Thea Whitman, Matthew F. Traxler

## Abstract

Wildfires represent a fundamental and profound disturbance in many ecosystems, and their frequency and severity are increasing in many regions of the world. Fire affects soil by removing carbon in the form of CO_2_ and transforming remaining surface carbon into pyrolyzed organic material (PyOM). Fires also generate substantial necromass at depths where the heat kills soil organisms but does not catalyze the formation of PyOM. *Pyronema* species strongly dominate soil fungal communities within weeks to months after fire. However, the carbon pool (i.e. necromass or PyOM) that fuels their rise in abundance is unknown. We used a *Pyronema domesticum* isolate from the catastrophic 2013 Rim Fire (CA, USA) to ask if *P. domesticum* is capable of metabolizing PyOM. *P. domesticum* grew readily on agar media where the sole carbon source was PyOM (specifically, pine wood PyOM produced at 750 °C). Using RNAseq, we investigated the response of *P. domesticum* to PyOM and observed a comprehensive induction of genes involved in the metabolism and mineralization of aromatic compounds, typical of those found in PyOM. Lastly, we used ^13^C-labeled 750 °C PyOM to demonstrate that *P. domesticum* is capable of mineralizing PyOM to CO_2_. Collectively, our results indicate a robust potential for *P. domesticum* to liberate carbon from PyOM in post-fire ecosystems and return it to the bioavailable carbon pool.

**IMPORTANCE:** Fires are increasing in frequency and severity in many regions across the world. Thus, it’s critically important to understand how our ecosystems respond to inform restoration and recovery efforts. Fire transforms the soil, removing many nutrients while leaving behind both nutritious necromass and complex pyrolyzed organic matter, which is often recalcitrant. Filamentous fungi of the genus *Pyronema* strongly dominate soil fungal communities soon after fire. While Pyronema are key pioneer species in post-fire environments, the nutrient source that fuels their rise in abundance is unknown. In this manuscript, we used a P. domesticum isolate from the catastrophic 2013 Rim Fire (CA, USA) to demonstrate that *P. domesticum* metabolizes pyrolyzed organic material, effectively liberating this complex pyrolyzed carbon and returning it to the bioavailable carbon pool. The success of Pyronema in post-fire ecosystems has the potential to kick-start growth of other organisms and influence the entire trajectory of post-fire recovery.

## INTRODUCTION

Wildfires can have substantial effects on nutrient cycling [1, 2] and community composition both above- and belowground [3, 4], making them important drivers of ecosystem processes [5]. Furthermore, wildfires are increasing in frequency and severity in many regions of the world [6]. Independent of soil type, wildfires have been shown to decrease the total amount of carbon in surface soils through combustion, releasing it as carbon dioxide, while much of the remaining carbon is transformed into black carbon, or pyrogenic organic matter (PyOM) [7–11]. PyOM encompasses a heterogeneous spectrum of compounds, but is predominantly composed of aromatic and polyaromatic compounds, depending on the source material, the temperature, and duration of pyrolysis [12–14]. PyOM is generally thought of as being relatively recalcitrant, with PyOM sometimes persisting for hundreds or thousands of years [9, 12]. While organic matter in surface soils may be completely combusted or pyrolyzed during fire, in deeper soil layers, non-pyrolyzed organic carbon is released where the heat from fire was enough to kill cells, forming a necromass zone, but not hot enough for combustion or to catalyze the formation of PyOM [8, 15]. Thus, post-fire soils often contain surface layers infused with PyOM, and necromass zones with abundant organic matter directly below. Early microbial colonizers of post-fire soils may exploit either or both PyOM and necromass as a key carbon source. However, relatively little is known about how the metabolism of these respective carbon sources may drive post-fire microbial succession and community recovery.

Many microorganisms are able to metabolize polyaromatic compounds with similarities to those found in PyOM, either completely or incompletely [16]. For example, white-rot fungi have been particularly well-studied for their ability to metabolize the phenolic polymer lignin. These fungi leverage a combination of peroxidases, laccases, and monooxygenases to initiate the degradation of lignin and other polyaromatic compounds [17–19]. Non-lignolytic fungi rely primarily on monooxygenases, especially cytochrome P450 monooxygenases, coupled with epoxide hydrolases to initiate the degradation of complex polyaromatic compounds [16, 19, 20]. Several common soil fungi have also been shown to degrade polyaromatic compounds [18]. These fungi include *Neurospora crassa*, which emerges from burned wood shortly after fire, and *Morchella conica*, which is a relative of pyrophilous *Morchella* species that often co-occur with *Pyronema* species [21–24].

Fruiting bodies of the genus *Pyronema* are among the first macrofungi to emerge from burned soil, doing so within weeks to months after fire [15, 24–26] (Figure 1 A&B). There are currently only two described species of *Pyronema*: *P. domesticum* and *P. omphalodes* (= *P. confluens*), both of which rapidly dominate post-fire fungal communities [15]. A recent ITS amplicon community analysis showed that *Pyronema* reads, which made up less than 1% of reads (0.91%) prior to fire achieved a post-fire average relative abundance of 60.34% [15]. Both *P. domesticum* and *P. omphalodes* were isolated from fruiting bodies that appeared within months after the catastrophic 2013 Rim Fire in Stanislaus National Forest, near the border with Yosemite National Park (California, USA) [15]. In vitro, *Pyronema* has a rapid growth rate, but has historically been considered a poor competitor with other soil fungi [27, 28]. Thus, a key question is: what carbon source is used by *Pyronema* to achieve such high relative abundance post-fire? Does *Pyronema* simply exploit the available necromass, or do they have the ability to metabolize PyOM as well? Given the dominant status and their early emergence after fire, *Pyronema* likely play a critical role in the first steps of post-fire succession. Thus, the possibility that *Pyronema* might contribute to the mineralization of PyOM has far-reaching implications for carbon cycling within post-fire soil communities.

**Figure 1:**
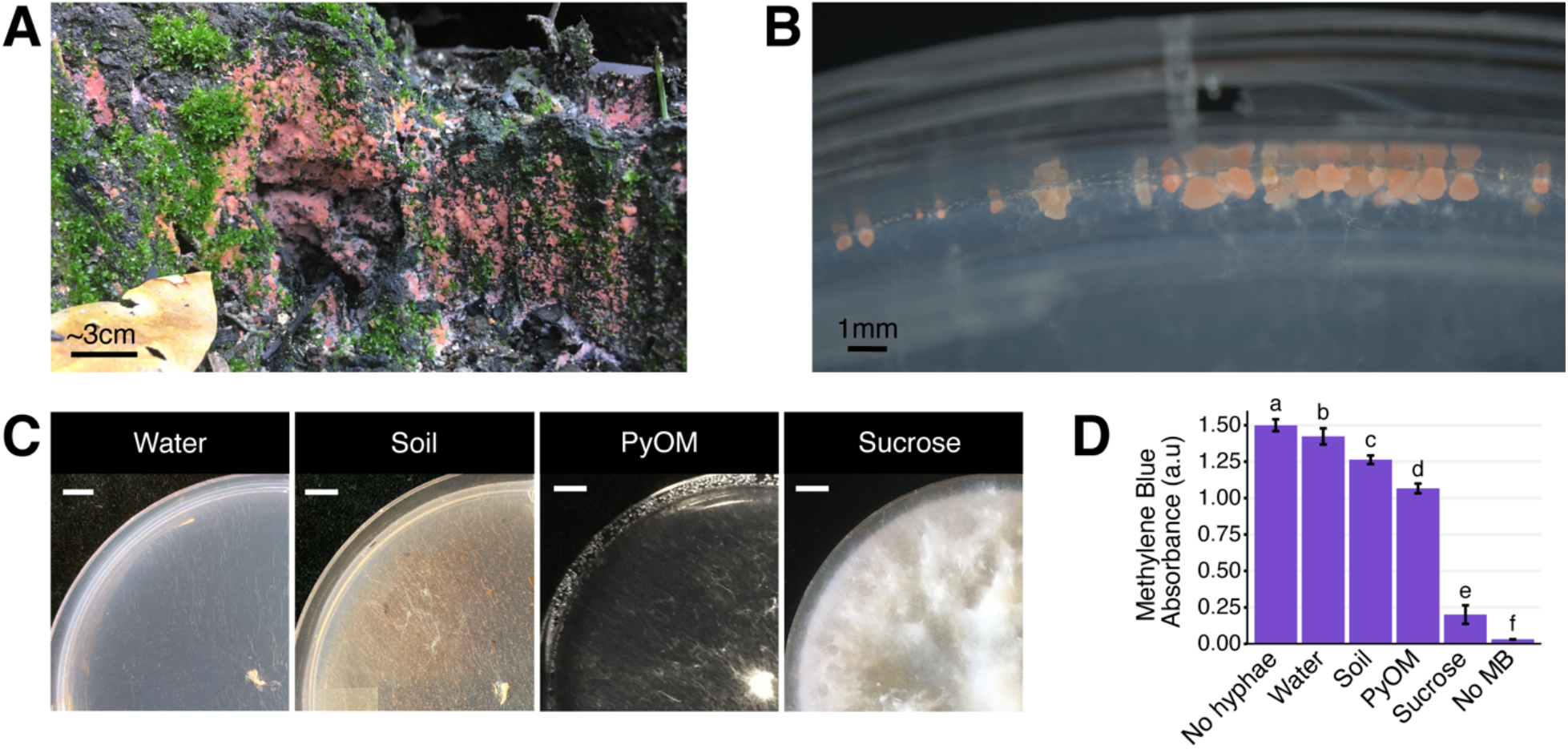
Pyronema growth on natural and laboratory substrates. (A) *Pyronema sp*. fruiting on burned soil six months after the 2018 Camp Fire began near Paradise, California, USA. (B) *Pyronema domesticum* DOB7353 ascocarps on the edge of a water agar plate. (C) *P. domesticum* DOB7353 growing on four different agar media treatments: water agar with no added nutrients (“Water”), wildfire-burned soil collected near the original isolation site for *P. domesticum* DOB7353 (“Soil”), 750 °C White Pine wood char (“PyOM”), or Vogel’s minimal medium with sucrose as a carbon source (“Sucrose”). All plates were inoculated in the center of the plate (bottom-right corner of photo) with mycelium from a 6mm-diameter punch of an actively growing *P. domesticum* colony. Scale bar = 1cm. (D) Average amount of *P. domesticum* DOB7353 biomass on one full plate as show in C, quantified by measuring the amount of Methylene Blue (MB) stain remaining after absorption by *P. domesticum* hyphae. All treatments are significantly different from each other (ANOVA + Tukey’s test, p < 0.0001, n = 5, error bars = standard deviation). The “No hyphae” control is pure 0.2mM MB without any biomass treatment. The “No MB” control is a blank well.

In this work, we investigated the hypothesis that early successional pyrophilous fungi such as *Pyronema* metabolize PyOM. To do so, we measured biomass, sequenced the transcriptome (RNAseq), and measured CO_2_ efflux from *P. domesticum* grown on agar media with various carbon sources, including PyOM and burned soil collected from a frequent and high-intensity wildfire site [29]. When grown on media containing burned soil or PyOM, *P. domesticum* produced significant biomass, activated a diverse suite of cytochrome P450 and FAD-dependent monooxygenases, and comprehensively induced pathways for aromatic substrate utilization. Lastly, we confirmed that *P. domesticum* mineralized PyOM by measuring CO_2_ emissions of *P. domesticum* grown on ^13^C-labeled PyOM. Collectively, our results demonstrate the potential for *P. domesticum* to liberate carbon from PyOM, assimilate it into biomass, and mineralize it to CO_2_. Thus, pioneering organisms such as *P. domesticum* may play an important role in the short-term reintegration of PyOM into biologically available carbon in post-fire ecosystems.

## MATERIALS AND METHODS

### Pyrogenic organic matter production

PyOM was produced from *Pinus strobus* (L.) (eastern white pine) wood chips <2 mm at 750 °C in a modified Fischer Scientific Lindberg/Blue M Moldatherm box furnace (Thermo Fisher Scientific, Waltham, MA, USA) fitted with an Omega CN9600 SERIES Autotune Temperature Controller (Omega Engineering Inc., Norwalk, CT, USA). We modified the furnace and adapted the PyOM production design developed by Güereña, *et al*. [30]. Briefly, the feedstock was placed in a steel cylinder inside the furnace chamber and subjected to a continuous argon gas supply at a rate of 1 L min^-1^ to maintain anaerobic conditions during pyrolysis. The heating rate for production of PyOM was kept constant at 5 °C min^-1^. We held the temperature constant for 30 min once 750 °C was reached, after which the PyOM was rapidly cooled by circulating cold water in stainless steel tubes wrapped around the steel cylinder. The PyOM was ground using a mortar and pestle and sieved to collect PyOM with particle size <45 µm.

### Fungal strain and biomass quantification

*Pyronema domesticum* DOB7353 [15] was inoculated onto 1.5% agar media treatment plates overlaid with cellophane; Vogel’s Minimal Medium [31] agar containing 20 g L^-1^ sucrose (“sucrose”), 10 g L^-1^ 750 °C PyOM agar (“PyOM”), 10 g L^-1^ wildfire-burned soil agar (“soil”), and water agar (“water”). Burned soil was collected from 0-10 cm in Illilouette Creek Basin [29] via an ethanol-sterilized shovel, and homogenized in plastic zip-top bags. Burned soil was x-ray sterilized (Steris, Petaluma, CA) and both PyOM and soil were added to agar media after autoclaving.

*P. domesticum* was allowed to grow for four days until it completely covered the plate on each agar media treatment described above (sucrose, PyOM, soil, and water). Biomass from each plate was harvested, immediately weighed, and then mixed with 500 μL 0.2 mM Methylene Blue (M9140, MilliporeSigma) in a 1.5 mL microcentrifuge tube. We adapted Fisher & Sawers’ Methylene Blue (MB) biomass quantification protocol [32]. Briefly, tubes of MB-stained biomass were heated at 80 °C for 5 minutes, then vortexed at maximum speed for 10 min, then heated again at 80 °C for 5 min. Mycelia was pelleted by centrifugation for 10 minutes at maximum speed in a standard microcentrifuge. 50 μL of the supernatant was combined with 200 μL ddH_2_O and then absorbance was measured at 660 nm. Blank wells, and wells containing 0.2 mM MB were included as controls.

### RNA extraction and sequencing

Mycelia was harvested from a total of nine replicate plates for each treatment (as described above). Mycelia from sets of three plates were pooled, resulting in three replicate samples for RNA extraction and sequencing. Pooled mycelia were immediately flash frozen with liquid nitrogen. Cells were lysed by bead-beating with 1 mL TRIzol [33]. Nucleosomes were removed by gently shaking for 5 minutes at room temperature. 200 uL chloroform was added, briefly bead-beaten, and then centrifuged to pellet cell debris. The aqueous phase was then used for RNA purification with the Zymo Direct-zol RNA MiniPrep kit (Cat. No. R2050). The qb3 facility at University of California, Berkeley quantified RNA quality and concentration via Bioanalyzer and then carried out library preparation and sequencing on an Illumina NovaSeq 6000 Platform.

### RNAseq data analysis

Raw reads were manually inspected for quality using FastQC v0.11.5, and then trimmed and quality filtered with Trimmomatic v0.36 [34]. HISAT2 v2.1.0 [35] mapped quality reads to the *P. domesticum* DOB7353 v1.0 genome [15, 36]. Raw counts per gene were generated with HTSeq v0.9.1 [37]. Raw counts were normalized, a PCA plot was generated, and differential expression was calculated with DESeq2 v1.24.0 on R v3.6.1 [38, 39]. To determine whether expression profiles were significantly different across treatments, we used PERMANOVA from the adonis() function from the vegan package v2.5-7 [40]. Functional gene annotations were downloaded from the Joint Genome Institute’s Mycocosm portal [36]. Additional annotation of specific genes was performed via protein-BLAST.

### ^13^C labeled PyOM and respiration experiment

^13^C-labelled 750 °C PyOM was produced from *Pinus strobus* as described above, except the biomass was from ^13^C-labelled seedlings. The ^13^C label was incorporated by pulse-labelling 2-year-old *P. strobus* seedlings with ^13^CO_2_, resulting in a δ^13^C value (relative to the standard vPDB) of +833.11%_0_ in the PyOM. We incubated the *P. domesticum* on ^13^C-labelled 750 °C PyOM agar (10 g L^-1^ PyOM) in 118.29 mL Mason jars, fitted with gas-tight lines, connected to an automated sample analyzer (“multiplexer”) that automatically samples the jar headspaces at regular intervals and quantifies the amount and isotopic signature of the headspace CO_2_ in a Picarro cavity ringdown spectrometer (multiplexer described in detail in Berry *et al*., *in review*). To conserve limited labelled material while maintaining moisture in the media, we layered 10 mL PyOM media over 30 mL water agar in the Mason jars. *P. domesticum* was inoculated using a punch from an identical ^13^C-labelled 750 °C PyOM agar plate. The jars were sealed and connected to the multiplexer, where they were measured every 48-72 hours for 57 days. Between measurements, jar headspace was flushed with a 20% O_2_, 80% N_2_, and 400 ppm CO_2_ gas mix designed to represent atmospheric conditions. Measurement frequency was such that jars did not become oxygen-depleted. We used five replicates of *P. domesticum*-inoculated plates and five replicates of control uninoculated plates.

CO_2_ emissions were partitioned between sources using stable isotope partitioning and the following equation [41]:

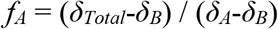

where *f*_*A*_ is the fraction of total CO_2_ emissions from source A, and *δ* represents the δ^13^C signature of the total (*δ*_*Total*_), source A (*δ*_*A*_), or source B (*δ*_*B*_). To calculate the CO_2_ that was released specifically due to the presence of *P. domesticum*, we subtracted the effects of abiotic sorption of CO_2_ by PyOM (red diamonds in Figure 5) from the total CO_2_ based on the emissions from the uninoculated jars, adjusting the isotopic signature accordingly. To determine the portion of the remaining biotic emissions that were derived specifically from PyOM, we then partitioned the remaining CO_2_ between PyOM and non-PyOM sources, using the δ^13^C value of the PyOM and the δ^13^C value of media-derived CO_2_ evolved from control, *P. domesticum*-inoculated water agar plates.

### Data Availability

We have provided an Excel file in the supplemental materials associated with the article, which details the results of our differential expression analysis and functional category assignment. FASTQ raw RNAseq data is publicly available at SRA accession PRJNA662999. Lastly, full code used for processing gas data is available at github.com/whitmanlab.

## RESULTS

### Pyrolyzed substrates induce a distinct transcriptional response

We observed distinct differences in the macroscopic growth pattern of *P. domesticum* when grown on four different agar media treatments; 750 °C *Pinus strobus* wood PyOM, wildfire burned soil, sucrose minimal medium, and water agar (Figure 1C). After inoculating agar treatment plates with equivalent amounts of mycelia, a substantial amount of biomass was produced on sucrose (Figure 1 C&D, and Figure S1). Growth on PyOM and, to a lesser extent, burned soil both produced an intermediate amount of biomass. Notably, *P. domesticum* has a tufted or fluffy macroscopic morphology on sucrose and to a lesser extent, PyOM. Lastly, there was observable growth on water agar, but biomass production was minimal (Figure 1 C&D, and Figure S1).

After four days of growth on each substrate, the biomass from each treatment was harvested, and RNA was extracted for sequencing. Principal Component Analysis (PCA) of these transcriptomes (Figure 2) illustrates the significant differences between treatments (PERMANOVA, p = 0.001, n=3). Across PC2 (23% of variation), the transcriptomes from the water and sucrose conditions fell at opposite ends, while transcriptomes from the PyOM and burned soil were located at an intermediate point near the origin. A possible explanation for this distribution is that PC2 describes the overall amount of bioavailable carbon and other nutrients. Water agar representing starvation contains the least amount of nutrients, the PyOM and soil containing intermediate amounts, and sucrose agar containing the most. Across PC1, which explained 56% of the variance across our samples, the PyOM -associated transcriptomes were located at one end of the axis while the water and sucrose conditions fell at the opposite end, with the burned soil transcriptomes at an intermediate position near the sucrose and water conditions. One possibility is that PC1 reflects the amount of PyOM present in the medium, since the PyOM medium contained the most, burned soil contained less, and sucrose and water media lacked any at all. Together, these results indicate that the transcriptional response of *P. domesticum* to burned or pyrolyzed substrates is unique compared to either water or sucrose, and the response to PyOM is particularly distinct.

**Figure 2.**
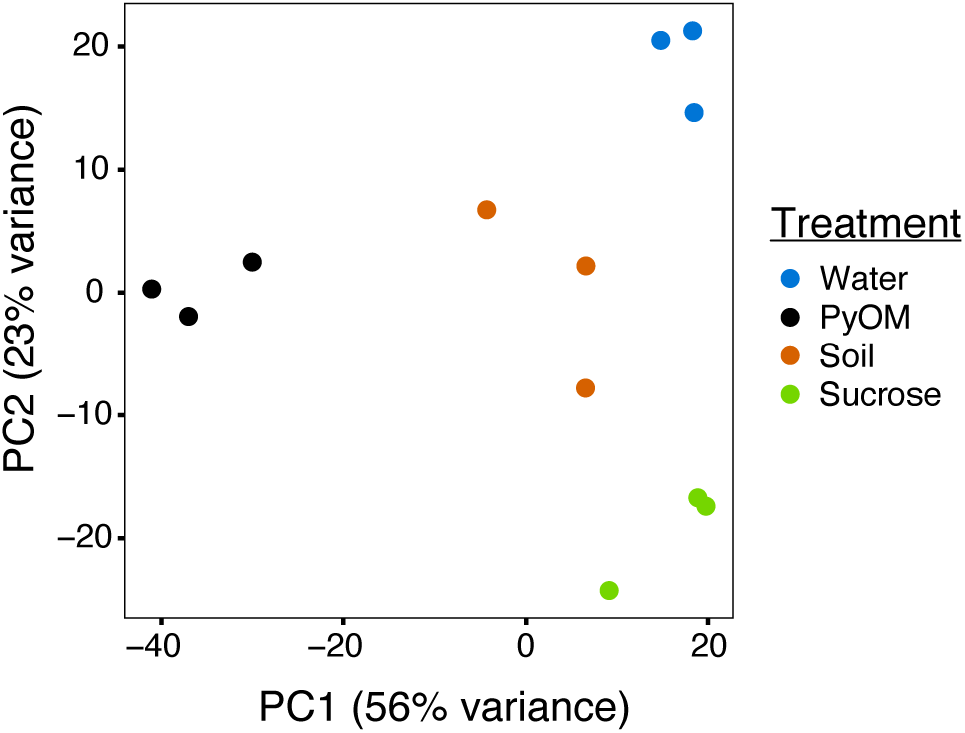
Pyrolyzed substrates induce expression of distinct sets of genes in *P. domesticum*. Principal Component Analysis plot illustrating the variation between each sample transcriptome (normalized expression values). Prior to RNA extraction, *P. domesticum* DOB7353 was grown in triplicate on four different agar media treatments.

### Starvation stress induces a broad transcriptional response

Growth on water agar triggered a broad starvation stress response in *P. domesticum* (Supplemental Data). Compared to sucrose, on water agar we observed significant upregulation of 318 genes (Figure 3A), including 31 transporters and 86 genes involved in the metabolism of diverse substrates, including the catabolism of amino acids and nucleotides (adjusted p-value < 0.01, fold change > 4, n = 3; Supplemental Data). Several general stress response genes were also induced on water agar compared to sucrose; specifically, seven different heat shock proteins and two proteins involved in programmed cell death. Surprisingly, invertase, the enzyme that hydrolyzes sucrose, was not significantly downregulated on water compared to sucrose (adjusted p-value = 0.14, fold change = 1.8, n = 3). In contrast to the 318 genes that were upregulated on water compared to sucrose, there were only 94 genes significantly upregulated on sucrose compared to water, including a sugar:hydrogen symporter, and 23 genes involved in primary metabolism, biosynthesis, and development (Supplemental Data). Taken together, these data demonstrate that growth on water agar induces a stress response program that includes genes involved in catabolism of macromolecules and scavenging for alternative nutrient sources. In contrast, growth on sucrose allows for a more streamlined transcriptome focused on growth powered by the metabolism of simple sugars.

**Figure 3.**
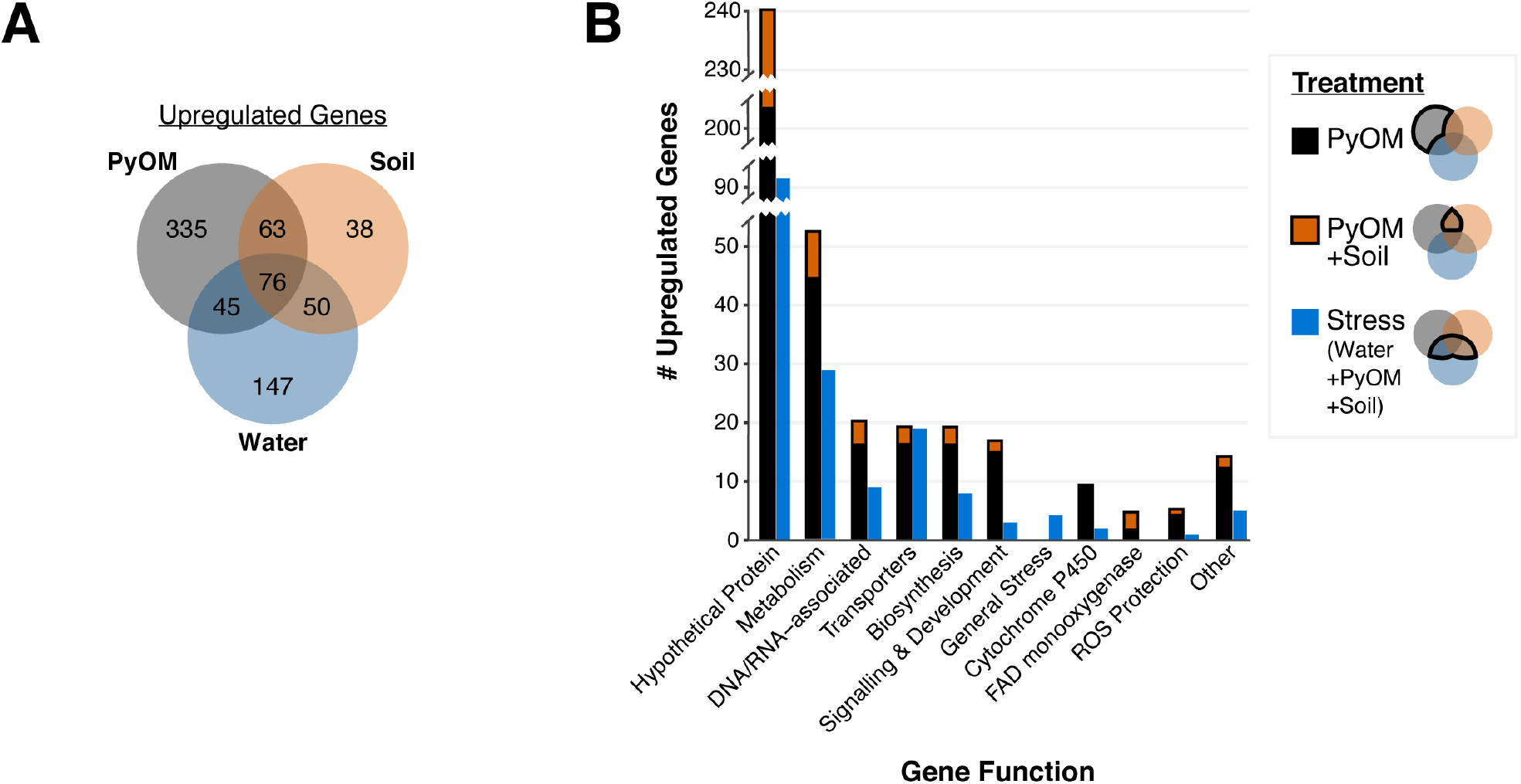
Pyrolyzed substrates induce expression of genes involved in stress response and PyOM metabolism. (A) Venn diagram showing the number of significantly upregulated genes in each treatment compared to sucrose (adjusted p-value < 0.01, fold change > 4, n = 3). (B) Number of significantly upregulated genes compared to expression on sucrose in each functional gene category (adjusted p-value < 0.01, fold change > 4, n = 3). Functional gene categories were determined via KEGG, GO, and pfam annotations. Stacked black and orange bars indicate the number of genes upregulated on PyOM alone (black) or the overlap between PyOM and soil (orange and black). We defined stress-response genes as those which are upregulated on water agar. Blue bars indicate the number of genes that are upregulated on both water and burned or pyrolyzed substrates for each functional category.

### The transcriptional response to pyrolyzed substrates is characterized by genes involved in stress tolerance, metabolism, and growth

To examine the nutritional and metabolic response to burned or pyrolyzed substrates, we calculated differential expression of genes in each treatment compared to sucrose and used functional gene annotations to categorize genes that were significantly upregulated at least 4-fold (Figure 3, for downregulated genes see Figure S2). We observed the largest shift in gene expression on PyOM with a total of 519 significantly upregulated genes (Figure 3A). 227 genes were upregulated on burned soil, and the majority (189 genes) of those overlapped with genes induced on PyOM and/or water (adjusted p-value < 0.01, fold change > 4, n = 3). We note that invertase was significantly down-regulated on PyOM compared to sucrose (adjusted p-value = 1.17E-7, fold change = -9.9, n = 3), and to a lesser extent on soil compared to sucrose (adjusted p-value = 0.02, fold change = -5.7, n = 3).

The 171 genes that were induced on water and at least one of the two substrates containing PyOM (burned soil and PyOM) characterized a stress response associated with decreased nutrient availability. Among these 171 genes are nineteen transporters and four general stress response genes including two heat shock proteins (Figure 3B, Supplemental Data). Additionally, we observed signatures of nitrogen stress in the water, PyOM, and soil conditions compared to sucrose minimal medium, which contains ammonium nitrate as a nitrogen source. These putative nitrogen stress responsive genes include genes involved in ammonium production, nitrogen metabolism, and a putative ortholog (gene_1304) of the conserved *Aspergillus nidulans* transcription factor TamA (Supplemental Data). TamA is a conserved stress-responsive regulator of nitrogen metabolism [42].

The 63 genes that were induced in common between PyOM and burned soil, excluding water, characterize a common response to PyOM (Figure 3). In addition, 335 genes were uniquely upregulated in response to PyOM, and the 38 genes uniquely upregulated on burned soil were almost entirely annotated as hypothetical proteins (Supplemental Data). After ‘hypothetical’, the next category with the most genes was that of metabolism, which we address in the subsequent section. We note that PyOM-responsive genes included nine Cytochrome P450 monooxygenases and four FAD monooxygenases. Cytochrome P450 oxidation of aromatic compounds often results in the formation of toxic epoxides and reactive oxygen species (ROS). On both substrates containing PyOM we observed upregulation of genes involved in ROS protection (Figure 3B). However, neither of the two epoxide hydrolases annotated in the *P. domesticum* genome exhibited any significant changes across our treatments (Supplemental Data). Lastly, we observed an enrichment of genes involved in biosynthesis (e.g., synthesis of amino acids, fatty acids, membrane lipids), development, and signaling that were upregulated specifically in the presence of PyOM. Taken together, these data indicate that, as expected, growth on PyOM is more stressful than growth on sucrose. Beyond a general stress response, the *P. domesticum* response to burned or pyrolyzed substrates includes the activation of a large set of genes, including those involved in metabolism, oxidation of aromatic substrates, and protection from ROS.

### PyOM induces a coherent set of metabolic pathways for aromatic compound degradation in P. domesticum

The results in the previous section indicate that PyOM may prompt a restructuring of metabolism in *P. domesticum*. In Figure 4 we mapped the significantly upregulated genes in *P. domesticum* (adjusted p-value < 0.01, fold change > 2, n = 3) onto the canonical pathways for aromatic compound degradation and assimilation into central metabolism and other biosynthetic pathways. All PyOM is enriched for aromatic carbon compounds because incomplete combustion of organic matter results in the formation of aromatic and polyaromatic carbon compounds [12]. PyOM produced at temperatures greater than ∼400°C generally has a carbon composition that is >90% aromatic [12, 43]. Here we propose that the large cohort of cytochrome P450 and FAD monooxygenases that were induced on PyOM-containing media (compared to growth on sucrose) are the primary method that *P. domesticum* uses to initiate the degradation of polyaromatic and aromatic carbon compounds. FAD monooxygenases oxidize compounds with a single aromatic ring, whereas cytochrome P450 monooxygenases can oxidize complex polyaromatic compounds [19, 44].

**Figure 4.**
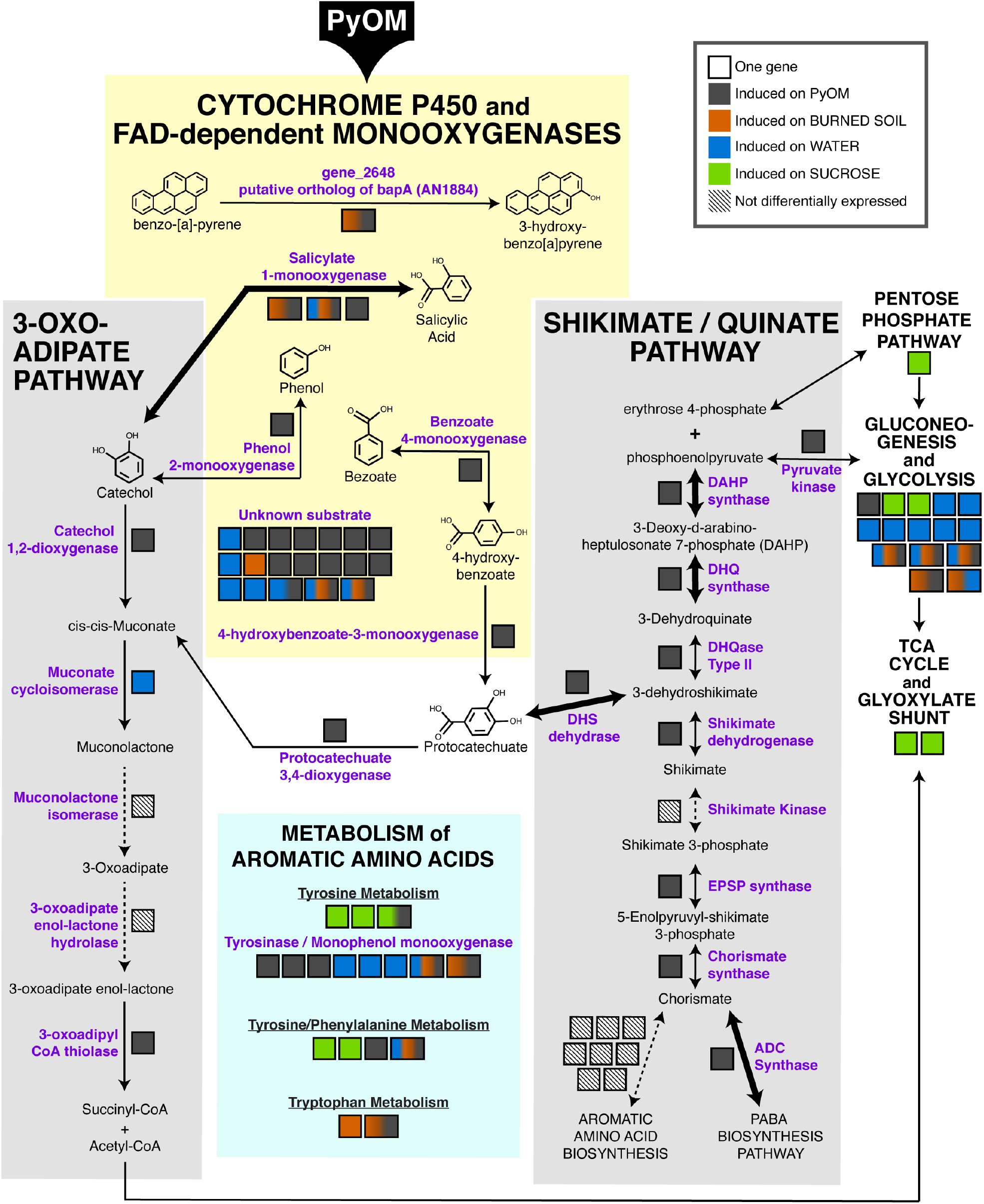
Metabolic map highlighting aromatic compound metabolism induced by growth on pyrolyzed substrates. Significantly upregulated genes mapped onto the canonical pathways for aromatic compound metabolism (adjusted p-value < 0.01, fold change > 2, n = 3). Bolded arrows indicate a fold change > 8 on PyOM compared to sucrose. Each gene is indicated as a black-outlined box, and the proteins encoded by these genes are indicated as purple text. The color fill of the box indicates the condition(s) in which the gene was upregulated. Multi-colored boxes are slightly larger than mono-color boxes to increase visibility of the colors and to highlight genes that are induced in more than one condition. Diagonal parallel lines within a box and associated dashed lines indicate genes that were expressed, but not differentially expressed under the tested conditions.

One cytochrome P450 gene (gene_2648) that was upregulated on both PyOM and burned soil was identified via protein-BLAST as a putative ortholog of the *bapA* gene in *A. nidulans*, which was recently shown to oxidize the polyaromatic hydrocarbon benzo-[a]-pyrene [19]. An additional five upregulated FAD monooxygenase genes and one cytochrome P450 monooxygenase gene have specific predicted substrates (salicylic acid, phenol, and benzoate). Lastly, fifteen cytochrome P450 monooxygenase genes were induced at least 2-fold on PyOM-containing media that have currently unknown substrates (Figure 4, Supplemental Table). Nearly half of these genes were strongly induced on PyOM; gene_10112, encoding a cytochrome P450 was strongly upregulated on both PyOM compared to sucrose (fold change = 1910.9) and on PyOM compared to water (fold change = 891.4), and six other cytochrome P450 genes were also upregulated at least 8-fold on PyOM compared to sucrose.

We identified two pathways by which aromatic carbon may be assimilated into central metabolism: via the protocatechuate and shikimate/quinate pathway and the via the catechol and 3-oxoadipate (=beta-ketoadipate) pathway. Six of seven core genes in the shikimate/quinate pathway were upregulated on PyOM compared to sucrose. In contrast, two of the five genes in the 3-oxoadipate pathway were upregulated on PyOM compared to sucrose. Notably, we observed strong upregulation on PyOM compared to sucrose of the four genes necessary to connect aromatic protocatechuate to central metabolism. These four genes encode DHS dehydrase (fold change = 36.8), DHQase (fold change = 4.6), DHQ synthase (fold change = 955.4), and DAHP synthase (fold change = 8.0). These three genes were similarly strongly upregulated on PyOM compared to water (Supplemental Data). In contrast, the genes that encode the proteins necessary for the 3-oxoadipate pathway were relatively modestly upregulated on PyOM compared to sucrose (fold change = 2.8, adjusted p-value < 0.01, n = 3).

We also observed that genes for the breakdown and metabolism of the three aromatic amino acids were induced differentially across all tested conditions. It is notable that upregulation of monophenol monooxygenase genes (i.e., tyrosinases) were also enriched on burned or pyrolyzed substrates and water compared to sucrose. Upregulation of central metabolism genes was generally enriched on sucrose, however some genes involved in glycolysis and gluconeogenesis were also upregulated on water, PyOM, and soil.

In summary, when *P. domesticum* was grown on PyOM, we observed upregulation of an extensive set of monooxygenases that may initiate degradation of the aromatic components of PyOM. We also observed comprehensive induction of the shikimate/quinate and 3-oxoadipate pathways, though the shikimate/quinate pathway was much more strongly induced. These data indicate that the aromatic intermediates liberated by monooxygenases may be funneled into central metabolism and mineralized via the shikimate/quinate and 3-oxoadipate pathways in *P. domesticum*.

### P. domesticum mineralizes PyOM-derived carbon to CO_2_

To conclusively determine whether *P. domesticum* was able to mineralize PyOM carbon, we cultivated it on agar plates supplemented with ^13^C-labelled 750 °C PyOM and quantified CO_2_ emissions (Figure 5). In the headspace of gas-tight jars inoculated with *P. domesticum* (n = 5), we observed a cumulative increase in the production of both total CO_2_ (Figure 5A) and ^13^C-labelled PyOM-derived CO_2_ (Figure 5B) over several days of observation. This result indicates that *P. domesticum* mineralized some of the PyOM by converting the carbon from 750 °C PyOM into CO_2_. We also observed an accumulation of non-^13^C-labelled CO_2_ in the inoculated jars, indicating that *P. domesticum* was also mineralizing carbon from non-PyOM sources. In uninoculated control jars, we observed a net sorption of CO_2_ by the PyOM agar medium. Taken together, these results indicate that mineralization of carbon from both PyOM and non-PyOM sources in the media by *P. domesticum* was greater than the effect of abiotic CO_2_ sorption by PyOM media.

**Figure 5.**
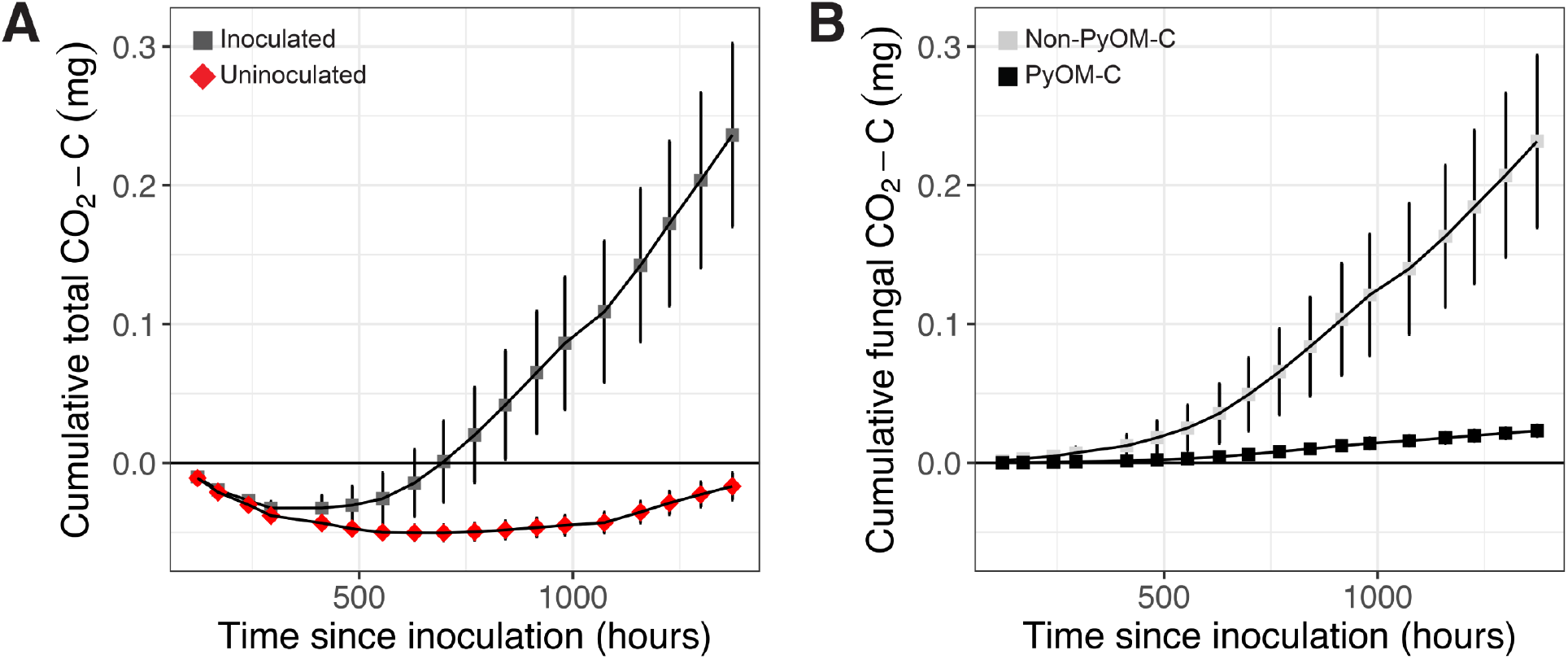
Cumulative mean CO_2_ emissions from *P. domesticum* growing on ^13^C-labeled 750°C PyOM. (A) Mean cumulative CO_2_ measured over time from the enclosed headspace of jars containing either sterile (uninoculated, red diamonds) 750 °C PyOM agar, or identical plates inoculated with *P. domesticum* (inoculated, dark grey squares) (n = 5, error bars = standard error). (B) Mean cumulative CO_2_ from *P. domesticum* inoculated jars, normalized by the uninoculated controls, and then partitioned into PyOM-derived C (black squares) and non-PyOM-derived C (light grey squares) using ^13^C partitioning (n = 5, error bars = standard error).

## DISCUSSION

Fungi in the genus *Pyronema* are pioneer species that rapidly dominate fungal communities in post-fire soils [15]. Thus, *Pyronema* have the potential to directly influence the trajectory of post-fire community succession and associated nutrient cycling dynamics. Here we investigated the transcriptional response of *Pyronema domesticum* on four different agar treatments: 750 °C PyOM, wildfire burned soil, sucrose minimal medium, and water. Our results indicate that burned or pyrolyzed substrates induce transcription of a comprehensive set of genes that together function to metabolize aromatic and polyaromatic compounds found in PyOM. Additionally, we demonstrated the mineralization of PyOM into CO_2_ by *P. domesticum*, consistent with the notion that this organism is capable of directly metabolizing PyOM.

*Pyronema* are barely detectable in soil before fire, become prevalent soon after fire, and then rapidly decline within weeks [15, 25]. The form taken by *Pyronema* between fire events is largely obscure. *Pyronema* may simply exist as dormant ascospores or sclerotia that require the heat and/or chemical changes associated with fire to trigger germination [27, 45]. One recent hypothesis suggests that pyrophilous fungi may live as endophytes for the majority of their life history, abandoning their plant hosts after they are killed by fire [46]. Regardless of how *Pyronema* live pre-fire, post-fire *Pyronema* are clearly poised to take full advantage of an open niche. Past work has shown that *Pyronema* are poor competitors, and they are also capable of growing rapidly on a diversity of substrates (i.e. burned soil, steam-treated soil, several different soil types, heat-treated plaster, and agar media containing various nutrients) [27, 28]. These data point toward the notion that *Pyronema* are generalists.

During intense forest fires, the organic material in the topmost layer of soil is heavily pyrolyzed, ultimately containing a significant amount of PyOM composed of complex aromatic and polyaromatic carbon compounds. A secondary layer of soil beneath the top layer is heated to a point that causes widespread death of the resident microbial/invertebrate soil fauna, leading to a layer rich in necromass that is not pyrolyzed. Carbon found in either layer could be targeted by *Pyronema*. The notion of *Pyronema* as generalists might suggest that they would be most likely to exploit the readily available carbon in the necromass layer. However, the metabolic restructuring at the transcriptional level, and production of ^13^C-labeled CO_2_ from labeled PyOM that we observed in this study, strongly indicate that *P. domesticum* readily metabolizes PyOM. Specifically, this restructuring includes the activation of an array of cytochromes P450 and FAD monooxygenases which likely target aromatic substrates for oxidation, in addition to activation of the shikimate/quinate and 3-oxoadipate pathways for assimilating the resulting substrates into central metabolic pathways. Thus, our results indicate that *Pyronema* may in fact be well-adapted as broad generalists able to capitalize on both necromass and abundant PyOM in post-fire soils, further explaining their rapid takeover of these communities.

Although their dominance is relatively short-lived in the post-fire community, *Pyronema* grow rapidly post-fire, producing abundant biomass in the form of ascocarps and mycelia [15, 24–26]. Competition may explain the short-lived dominance of *Pyronema* as it appears to be a weak competitor in isolation [28]. Even if *Pyronema* are outcompeted and simply senesce, their DNA could linger in post-fire soil and continue to be detected via sequencing methods [47–49]. However, both *Pyronema* ascocarps and their DNA decline rapidly after they peak in abundance following fire [15]. This rapid decline of *Pyronema* DNA could be explained by the starvation response that we observed on water agar (Figure 3), in which *P. domesticum* may fuel outward expansion perceived as growth by recycling macromolecular building blocks such as nucleotides and amino acids into a diffuse biomass aimed at exploration of environments with sparse nutrients [50, 51]. This turn-over of biomass may explain the non-PyOM-derived CO_2_ mineralized by *P. domesticum* (Figure 5B). Alternatively, *P. domesticum* may simply be mineralizing other carbon sources that were present in the agar medium, such as the agar itself. To our knowledge, terrestrial fungi lack agarases that degrade agarose, but the genomes fungi such as *P. domesticum* do contain a suite of pectinases, some of which may target agaropectin [52].

Another explanation for the rapid decline of *Pyronema* DNA in post-fire soils is that *Pyronema* biomass, either living or recently senesced, is consumed by other organisms [47, 48]. Thus, abundant *Pyronema* biomass may provide a critical nutrient source for secondary colonizers of post-fire soils, thereby laying the foundation for succession within post-fire communities. Importantly, the ability of *P. domesticum* to convert PyOM into biomass could directly facilitate the growth of organisms that lack the ability to metabolize PyOM. Thus, *Pyronema* may provide an important mechanism for rapidly assimilating some portion of newly formed PyOM back into more readily bioavailable forms of carbon in post-fire environments. Additionally, it is possible that *Pyronema* function to favorably transform the post-fire soil environment in other ways, such as affecting pH or accessibility of other nutrients. Nevertheless, the mineralization of PyOM by the dominant early-successional fungus *P. domesticum* is likely to have broad impacts on post-fire succession and recovery in soil microbial communities.

## Acknowledgements

We are grateful for Thomas D. Bruns who provided the *P. domesticum* strain and fruitful discussions. We thank Scott Stephens and Katya Rakhmatulina for generously allowing us to collect burned soil from their study sites in Illilouette Creek Basin at Yosemite National Park. This research was supported by the DOE Office of Science, Office of Biological and Environmental Research (BER), grant no. DE-SC0020351 to T.W. and M.F.T. The authors of this manuscript have no competing interests to disclose.

## SUPPLEMENTAL FIGURES

**Supplemental Figure 1:**
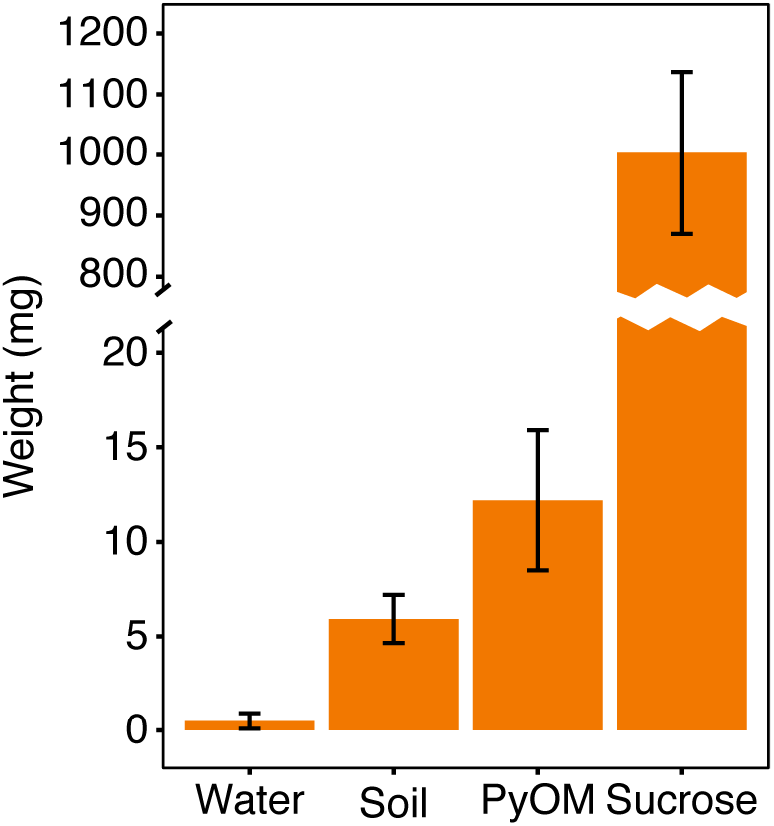
*P. domesticum* biomass wet weight on agar media treatments. Prior to Methylene Blue staining, *P. domesticum* biomass was harvested and immediately weighed. Error bars indicate standard deviation (n = 5).

**Supplemental Figure 2.**
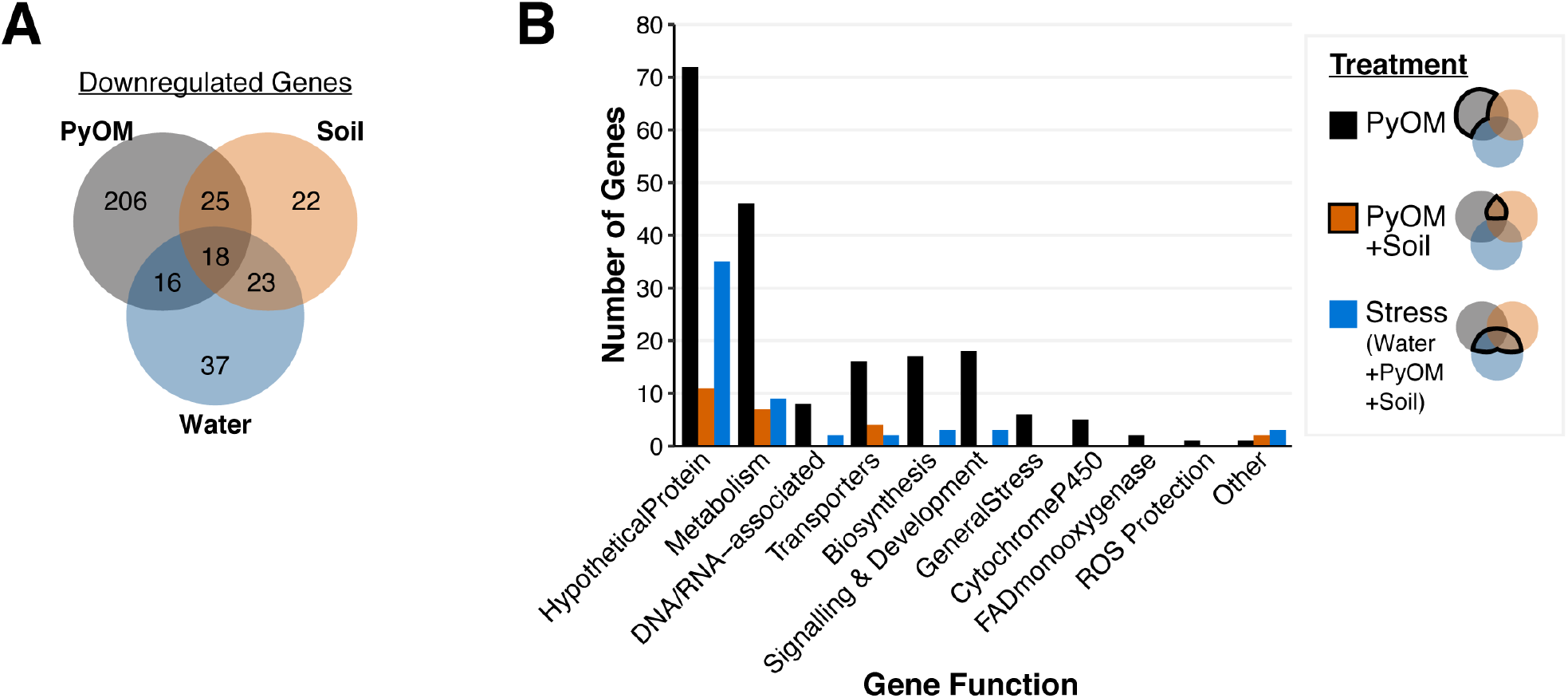
Down-regulated genes on pyrolyzed substrates in *P. domesticum*. (A) Venn diagram showing the number of significantly downregulated genes in each treatment compared to sucrose (adjusted p-value < 0.01, fold change < -4, n = 3). (B) Number of significantly downregulated genes compared to expression on sucrose in each functional gene category (adjusted p-value < 0.01, fold change < -4, n = 3). Functional gene categories were determined via KEGG, GO, and pfam annotations. Black bars indicate the number of genes downregulated on PyOM alone (total = 206). Orange bars indicate the number of genes downregulated in both PyOM and soil (total = 25). Blue bars indicate the number of genes downregulated on water that also overlap with soil and/or PyOM (total = 57).

